# Phylogenomics of *Plasmopara halstedii* reveals genomic regions associated with the breakdown of sunflower downy mildew resistance genes

**DOI:** 10.1101/2025.06.18.660304

**Authors:** Yann Pecrix, Etienne Dvorak, Frédéric Labbé, Ludovic Legrand, Sebastien Carrère, Jérôme Gouzy, François Delmotte, Guillaume Besnard, Laurence Godiard

**Affiliations:** Laboratoire des Interactions Plantes-Microbes Environnement (LIPME), INRAE, CNRS, Université de Toulouse, 31326 Castanet-Tolosan, France; CIRAD, UMR PVBMT, Saint Pierre, France; INRAE, Université de Bordeaux, Bordeaux Sciences Agro, SAVE, ISVV, F-33140, Villenave d’Ornon, France; Centre de Recherche sur la Biodiversité et l’Environnement (CRBE), Université de Toulouse, CNRS, IRD, Toulouse INP, Université Toulouse 3 – Paul Sabatier (UT3), Toulouse, France

**Keywords:** Evolutionary history, comparative genomics, resistance breakdown, admixture mapping, oomycete plant pathogen

## Abstract

- Understanding the genetic diversity and evolutionary history of plant pathogens is crucial for effective disease management strategies. Sunflower downy mildew, caused by the oomycete *Plasmopara halstedii*, is a worldwide threat to the sunflower oil crop. We aimed to explain through phylogenomic studies how downy mildew resistance breakdown occurred recurrently in the last decades in France, leading to new virulence profiles.
- We assembled high-quality genomes of three founder pathotypes of *Pl. halstedii*. Performing comparative genomic analyses, population genetics, and phylogenomic analyses, we studied the genomic structure among the 16 reference French pathotypes of *Pl. halstedii*.
- We revealed a conserved genomic organisation among pathotypes and a strong synteny with other Peronosporales species. The history of *Pl. halstedii* invasion in France over the last 60 years was documented by identifying founder strains and their admixture patterns. The emergence of pathotypes with broader virulence spectra and therefore capable of overcoming host resistance was associated with genomic reshuffling. We highlighted genomic mosaicism in admixed pathotypes and identified regions associated with the breakdown of host resistance genes harbouring putative effector genes.
- Our findings provide insights into evolutionary mechanisms underlying plant pathogen host adaptation, which has implications for a sustainable deployment of multiple resistance genes.

## INTRODUCTION

Disease genetic resistance is one of the main pathways for controlling plant pathogens (Thomas *et al*., 2024). A major concern for sustainable agriculture is to preserve their effectiveness over time (Coomber *et al*., 2024), which requires a good knowledge of the biology and evolution of pathogens. The adaptation of pathogens to plant resistance genes indeed depends on a number of factors and several mechanisms are involved in the emergence of new virulent strains. First, the efficacy of natural selection depends on the effective population size of pathogens and also on their ability to recombine. Second, their reproductive mode (sexual, clonal, or both) will thus strongly impact their evolutionary potential to develop new adaptations to their host and environment (Zhan *et al*., 2007). In particular, the admixture (hybridization) between strains could lead to better adapted pathotypes that combine, for instance, different avirulence genes (Man in ‘t Veld *et al*., 2007; Ahmed *et al*., 2012). Third, the pathogen dispersal, usually facilitated by agricultural trade and transport of infected plant material, is another key trait by impacting the probability of contact between strains. Finally, at the genomic level, there is also increasing evidence suggesting that single mutations and structural variations play an important role in the evolutionary processes leading to resistance breakdown (Hartmann, 2022). Following the two-speed structure of pathogen genomes, avirulence genes have been described to be associated to the dispensable part of the genome which is largely enriched in transposable elements acting as drivers of structural variations (Raffaele & Kamoun, 2012; Dong *et al*., 2016). The identification of avirulence genes and a deep knowledge of the proximal genomic mechanisms that underlie adaptation of the pathogen is therefore needed to reach durable management of plant resistance.

The oomycete *Plasmopara halstedii* (Farl.) Berl. & de Toni is an obligate parasite causing downy mildew disease on annual *Helianthus* species such as sunflower, an economically important crop, especially in France and eastern Europe (Spring, 2019). *Pl. halstedii* is diploid and homothallic, i.e. the same organism produces both sexual reproductive cells (Spring, 2000). Asexual generations occur during the plant growing season but sexual reproduction produces overwintering oospores (Spring and Zipper, 2000). Severe symptoms in sunflower (i.e. plant dwarfism, leaf bleaching, sporulation, and production of infertile flowers) strongly impair seed yield (Gascuel *et al*., 2015). The most common method for disease control is the introduction of resistant genes introgressed from wild sunflower species into cultivated varieties. However, the use of a few resistance sources was overcome by *Pl. halstedii*, leading to the emergence of new pathotypes whose origins remain unclear. *Plasmopara halstedii* is assumed to have originated in the central portion of the North American continent (Leppik, 1962), where it infects wild *Helianthus* species. It is generally accepted that this pathogen was introduced into Europe at the beginning of the XX^th^ century with the importation of sunflower seeds during early cultivation of commercial sunflower in the former USSR. *Plasmopara halstedii* has then shown a strong diversification in Europe, especially during the last 50 years (Ahmed *et al*., 2012). In France, epidemiological data of the disease and accurate inventories of the different pathotypes have been recorded by Terres Inovia (formerly CETIOM; https://www.terresinovia.fr) since the 60’s. More than ten *Pl. halstedii* pathotypes, which are classified by their differential virulence profiles in a set of sunflower inbred resistant lines, have indeed emerged in France in the last three decades (Gascuel *et al*., 2015). At least six of them have been recorded worldwide (Viranyi *et al*., 2015). The new pathotypes are responsible for the breakdown of most of the resistance loci used in current breeding programs of sunflower (Moinard *et al*., 2006; Tourvieille *et al*., 2012). To avoid breakdown of resistance, breeders favour the stacking of resistance genes into sunflower genotypes. Twenty-two major resistance genes, denoted *Pl*, have been reported and mapped in sunflower (Qi *et al*., 2016; Pecrix *et al*., 2018a; Molinero-Ruiz, 2022). Among them, ten new resistance factors discovered from wild *Helianthus* species and wild *Helianthus annuus* ecotypes (designated *Pl23–Pl32*) are effective against at least 16 downy mildew pathotypes (Pecrix *et al*., 2018a). It is important for cultivated sunflower varieties to carry as wide a range of resistance genes as possible, since over the past 50 years, the presence of only a few of them in most sunflower varieties led to the appearance of new virulent pathotypes of *Pl. halstedii* (Ahmed *et al*., 2012; Gascuel *et al*., 2015).

The recurrent resistance gene breakdown observed in field conditions resulted from the rapid evolution of effector genes of the pathogen (Gascuel *et al*., 2016a). Effectors are secreted pathogenicity factors which are required for the developmental cycle of plant-pathogenic oomycetes, in particular by altering host defence reactions and advancing the infection process (Fawke *et al*., 2015). During sunflower infection, *Pl. halstedii* shows intercellular growth, produces haustoria in plant cells, and expresses RxLR and CRN–type effectors, as many other oomycetes (As-sadi *et al*., 2011; Gascuel *et al*., 2015, 2016a; Sharma *et al*., 2015; Mestre *et al*., 2016; Wang *et al*., 2023). *Plasmopara halstedii* also expresses putative secreted effectors harbouring the WY domain fold, a structural domain found in many Peronosporales effectors associated or not with a RxLR motif (Boutemy *et al*., 2011). Thirty putative « core » RxLR effectors conserved in 17 *Pl. halstedii* pathotypes were previously identified by an effectoromics approach (Pecrix *et al*., 2019). Their transient expression in sunflower leaves revealed a wide diversity of targeted subcellular compartments, including chloroplasts and processing bodies. More than half of the 30 core effectors were putative true virulence factors of *Pl. halstedii* because they were able to suppress pattern-triggered immunity in *Nicotiana benthamiana*, and five of them induced hypersensitive responses in sunflower broad-spectrum resistant lines (Pecrix *et al*., 2019). The hypersensitive response triggered by the core effector PhRxLR-C01 co-segregated with the *Pl22* resistance, establishing a gene-for-gene interaction between sunflower and its pathogen (Pecrix *et al*., 2019).

Previous studies performed on 146 samples, representing the 14 French pathotypes recorded at the time, indicated three distinct groups, focused on the three prevalent pathotypes 100, 703, and 710 (Delmotte *et al*., 2008; Ahmed *et al*., 2012). Pathotype 100, the first one to be challenged by sunflower resistance gene *Pl1*, is no longer recorded in France, while a very close and more virulent pathotype (304) has emerged. These studies have also demonstrated that multiple introductions of *Pl. halstedii* have aided its establishment in France, and suggests that pathotypes co-occurrence facilitated their recombination thus driving the emergence of new and endemic pathotypes in response to host resistance. A comprehensive knowledge of the molecular mechanisms underlying the evolution of pathogenicity of *Pl. halstedii* is currently the most important issue for a sustainable control of sunflower downy mildew in the field. This requires the completion of high-quality reference genomes obtained by long-range sequencing methods, and as a prerequisite, the production of high molecular weight genomic DNA from spores (Penouilh-Suzette *et al*., 2020). Resequencing of the whole-genome of French *Pl. halstedii* pathotypes makes it possible to go far beyond traditional population genetics analyses and identify large genomic rearrangements, local genome variations (e.g. gene duplication or deletion), and single nucleotide polymorphisms (SNP). Such SNPs can also be used to identify genetic clusters (e.g. initially introduced pathotypes) and detect possible admixture (Delmotte *et al*., 2008). Finally, such genomic studies coupled with geographic and chronologic data of pathotypes appearance should help to retrace the recent history of *Pl.halstedii*.

Here, we aimed to document the history of introduction and subsequent diversification of *Pl. halstedii* in France, and to assess the extent of recombination in new pathotypes in relation to their virulence profiles. We used the PacBio technology to sequence the genome of three pathotypes (#304, 703, and 710), representing the three main lineages that were historically introduced into France (Ahmed *et al*., 2012; Gascuel *et al*., 2016a). We generated, for the first time, full length (ca. 72 to 75 Mb), high-quality reference genome sequences for the three pathotypes. Illumina genome sequences of 14 additional *Pl. halstedii* pathotypes (Pecrix *et al*., 2019), mainly from France, were then compared to these three reference genomes to analyse DNA polymorphisms. The history of *Pl. halstedii* invasion in France was inferred from the genomic structure of these 17 pathotypes, the timeline of their emergence, and their virulence profiles on differential sunflower lines. Finally, admixture blocks between the three PacBio sequenced pathotypes were localised, together with 131 putative secreted effectors, since the evolutionary fate of such pathogenic factors could give insights into breakdown mechanisms of sunflower resistance to *Pl. halstedii*.

## MATERIAL AND METHODS

### Sampling of *Plasmopara halstedii* pathotypes

Seventeen *Pl. halstedii* pathotypes, including all 16 reference pathotypes present in France (namely 100, 300, 304–10 (hereafter 304), 304–30, 307, 314, 334, 700, 703, 704, 707, 710, 714, 717, 730, and 774) and one pathotype from Spain (330), were previously collected by Tourvieille *et al*. (2012) and all sequenced with Illumina short reads (Pecrix *et al*., 2019). In this study, we selected the three isolates 304, 703, and 710 for long-read sequencing. They correspond to non-admixed pathotypes and are considered representative of the *Pl. halstedii* diversity presently found in France (Delmotte *et al*., 2008; Tourvieille *et al*., 2012). Spore suspensions samples were collected by gently scraping the surface of infected cotyledons, pelleted by centrifugation at 1,000 g for 1 min, frozen in liquid nitrogen and mechanically ground with beads using a Mixer Mill MM 400 (Retsch, Haan, Germany). High molecular weight genomic DNAs were extracted from spores as described by Penouilh-Suzette *et al*. (2020).

### Genome sequencing and assembly of three pathotypes

A genome of the 710 pathotype was previously released based on short reads (Pecrix *et al*., 2018a). Here, a meta-assembly strategy based on PacBio RSII data was applied to assemble the 304, 703, and 710 genomes, using the approach described for the *Medicago truncatula* genome (Pecrix *et al.,* 2018b). Table S1 provides details of the steps of the process including data, software, and the evolution of the assembly metrics. The meta-assembly started by a primary assembly performed with CANU v.1.8 (Koren *et al*., 2017). The corrected reads generated during the CANU assembly were also used as input of SMARTdenovo (https://github.com/ruanjue/smartdenovo), wtdbg2.0, and wtdbg2.3 (Ruan & Li, 2020). Then, contig sequences of these four primary assemblies were transformed into pseudo long reads of 100 kb with an overlap of 50 kb, which were merged and assembled with CANU. Two iterations of polishing were performed on the meta-assembly output using the quiver PacBio software, followed by two additional iterations of polishing using Illumina paired-end reads (Pecrix *et al*., 2019) and pilon v.1.22 (Walker *et al*., 2014). The taxonomic origin of all contigs was determined using several contig-level features, i.e. GC percent, spanning coverage (with both Illumina and PacBio reads), repeat content, high scoring taxonomy group, and absence of a significant similarity with the two other genomes (Table S2; Pecrix et al., 2019). We thus identified likely contaminant sequences and removed them (i.e. for a total of 0.036 Mb, 0.253 Mb, and 14.6 Mb for 703, 304, and 710 assemblies, respectively). For metrics comparisons, four *Pl. halstedii* genome assemblies were retrieved from the NCBI database (on 12/15/2022): OS-Ph8-99-BlA4 (Sharma *et al*., 2015), A23 Pilon (assembly GCA_004380875.1), A23R734 (assembly GCA_003640465.1), and the first draft genome of 710 (Pecrix *et al*., 2018a). The assembly metrics were obtained using QUAST v.5.0.0 (Gurevich *et al.,* 2013), BUSCO v.5.2.2 (Manni *et al*., 2021), and EukCC v.2.1.0 (Saary *et al*., 2020).

### Prediction of gene models

Protein and non-protein coding gene models were predicted using the integrative EuGene pipeline release 1.5 (Sallet *et al*., 2019; http://eugene.toulouse.inra.fr/Downloads/egnep-Linux-x86_64.1.5.tar.gz). Two protein databases were used with NCBI-BLASTx to contribute to translated regions detection: Swiss-Prot (November 2017) and the oomycete subset of GenBank (February 2018). Proteins similar to REPBASE 20.05 (Bao *et al*., 2015) were removed from the two datasets to avoid the integration of TE-related proteins in the training steps. The cleaned datasets were aligned with NCBI-BLASTX on the genomes. Chained alignments spanning less than 80% of the length of the database protein were removed. Transcriptome datasets of *Pl. halstedii* generated in previous studies (As-Sadi *et al*., 2011; Mestre *et al*., 2015; Pecrix *et al*., 2018a) were integrated as transcript evidence. RNAseq reads were assembled with velvet (Zerbino & Birney, 2008). The longest open reading frame regions were extracted from transcript contigs (min. 60 bp) and used as transcriptional evidence by the EuGene pipeline. Spliced alignments spanning at least 30% of the transcript length at a minimum of 97% identity were retained. Regions spanned by transcript alignments were preserved from the repeat masking process integrated in the pipeline. The same annotation protocol was used for the three genomes.

### Functional annotation of protein-coding genes

Protein coding genes of the three genomes were functionally annotated by integrating a BLASTp search of reciprocal best hits with 53 *Pelanosporum* proteins tagged as “reviewed” in the Uniprot database (90% span, 80% identity) as of February 2019 (Uniprot Consortium, 2013) and protein domain annotation using the Interpro database r.72.0 (Finn *et al*., 2017). The protein annotations were validated by the tbl2asn software (https://www.ncbi.nlm.nih.gov/genbank/tbl2asn2, February 2019). GO terms were assigned using the BLAST2GO pro software integrating BLASTp similarities with GenBank NR database (Benson *et al*., 2013) and Interpro r72.0 results.

### Variant calling

The *Pl. halstedii* 710 genome (release Plhal710_r2, this work) was used as the reference genome. The Illumina datasets (Table S3) were mapped with bwa v.0.7.15-r1140 (Li & Durbin, 2010). Several successive steps of filtering and merging (view -f 0×02, fixmate -c –r, merge, rmdup, and view -q 1 -F 4 -F 256) were performed using samtools v.1.3.1 (Li *et al*., 2009). A per pathotype variant calling was performed with samtools (mpileup –B) and varscan v.2.4.3 (Koboldt *et al*., 2012) (mpileup2snp --min-coverage 20 --min-reads2 10 --min-avg-qual 15 --min-var-freq 0.2 --min-var-freq-for-hom 0.75 --*p*-value 0.01 --output-vcf 1). Then, the complete list of polymorphic positions in at least one sample were collected and used to generate the global VCF file (samtools mpileup -B -l polymorphic_positions.txt, varscan mpileup2cns --min-coverage 20 --min-reads2 10 --min-avg-qual 15 --min-var-freq 0.2 --min-var-freq-for-hom 0.75 --*p*-value 0.01 --output-vcf 1). Finally, the VCF file was annotated using SnpEff v.4.3t (Cingolani *et al*., 2012) (eff -upDownStreamLen 1000 -no-downstream -no-intron -no-intergenic).

### Secretome and effector predictions of the three completed *Pl. halstedii* genomes

In the three genome assemblies, we predicted putative secreted proteins using SignalP v.5.0 (Almagro Armenteros *et al*., 2019) and then performed *in silico* genome-wide identification of *Pl. halstedii* RxLR effectors using HMM and heuristic approaches (Bhattacharjee *et al*., 2006; Whisson *et al*., 2007; Win *et al*., 2012). We did an independent search to add as potential effectors the secreted proteins having at least one WY-domain fold (Win *et al*., 2012).

### Genetic diversity and population structure

Population structure was determined using the parametric Bayesian model-based clustering method implemented in STRUCTURE v.2.3.4 (Pritchard *et al*., 2000). The analysis was performed for a number of clusters (*K*) ranging from 2 to 10. For each *K* value, runs were performed applying 100,000 Markov Chain Monte Carlo (MCMC) repetitions and a 10,000 burn-in period. In addition, we used a discriminant analysis of principal components (DAPC) implemented in the adegenet R package v.2.1.10 (Jombart *et al*., 2010).

### Comparative genomics

Syntenic blocks between the three genome assemblies were identified and visualised with the MCscan tool implemented in JCVI (Tang *et al*., 2024). Orthologous CDS were identified between assemblies using a C-score cut-off of 0.99 to find reciprocal best hits. The Plhal710_r2 assembly was used as the reference for visualisation because of its higher contiguity. Synteny between the three *Pl. halstedii* genomes and the *Peronospora effusa* chromosome-scale assembly (Fletcher *et al*., 2022) was assessed using SynMap v.2.0 (Haug-Baltzell *et al*., 2017). Contigs of the *Pl. halstedii* genomes were ordered and oriented according to the syntenic path generated by SynMap. The positions of matching CDS were found using the LAST algorithm (standard parameters) and were retrieved to be plotted as links. Plots were generated using Circos v.0.69.9 (Krzywinski *et al*., 2009).

### Analysis of pathotype admixture using genome-wide local ancestry

A phylogenomic approach was finally used to investigate genome-wide local ancestry among French pathotypes, considering four reference genomes (i.e. 100, 334, 703, and 710) as putative ancestral donors (see below). Admixture was thus analysed individually for 12 pathotypes (i.e. 704, 314, 714, 717, 300, 304, 304 −30, 307, 707, 700, 730, and 774). The dataset was filtered by keeping only variants of the highest quality, i.e. we filtered out any variant having a proportion of missing genotypes higher than 20%, and a minor allele frequency (MAF) less than or equal to 1% using VCFtools v.0.1.16 (Danececk *et al*., 2011), resulting in a total of 92,570 bi-allelic variants. To allow comparison at the chromosome scale, contigs were ordered and oriented based on the *Pe. effusa* genome using the syntenic path generated by SynMap. The genomic coordinates of *Pl. halstedii* variants were converted to the positions of the pseudo-assembly using a custom python script (https://github.com/fredericlabbe/Phalstedii_Phylogenomics). For each 100-kb non-overlapping window along the chromosomes, we computed a maximum likelihood (ML) tree using PhyML v.3.3.3 (Guindon *et al*., 2009) with a GTR substitution model and 100 bootstrap replicates. PhyML was run using script phyml_sliding_windows.py (https://github.com/simonhmartin/genomics_general/blob/master/phylo) with a minimum of 10 sites per window and a minimum of 10 non-missing genotypes per individual per window. As there are 15 possible topologies for an unrooted five-taxon tree (Fig. S3), we used TWISST to estimate the weight (frequency) of each topology among all the 751 non-overlapping windows across the genome (Martin & van Belleghem, 2017). We investigated genetic polymorphic variation associated with *Pl* resistant gene breakdown, by only considering homozygous and biallelic SNPs with a perfect distribution of both alleles between bulks of virulent and avirulent pathotypes.

## RESULTS

### Genome features and secretome of three historical French pathotypes of *Pl. halstedii*

We sequenced and annotated the genome of three *Pl. halstedii* pathotypes: 304, 703, and 710. The size of these three assemblies ranges from 72.1 to 74.8 Mb (Table 1), which is similar to the average genome length for oomycetes (74.9 Mb; McGowan & Fitzpatrick, 2020). The seven available *Pl. halstedii* assemblies have similar lengths, although, for pathotype 710, the new version presented an increased size, from 67.6 to 74.8 Mb (Plhal710_r2; Table 1). This comparison also revealed that the three new assemblies are devoid of ‘N’ and are more contiguous than the four GenBank genomes as they have higher N50 values and are composed of less contigs (i.e. ranging from 88 to 122 compared to the previous assemblies that contained 745 to 7,628 scaffolds). These assemblies display high levels of completeness since we found at least 95.2% of conserved genes from the Stramenopiles and Alveolata databases at the genome level. Repetitive elements make up approximately half (49%) of the genomes, most of them (81%) being Long Terminal Repeat (LTR) retrotransposons. This high representation of LTRs in the repeated sequences is shared with *Pl. viticola* (79%; Dussert *el al.*, 2019) and *Pe. effusa* (74%; Fletcher *et al*., 2022).

**Table 1.** Compared characteristics of *Plasmopara halstedii* genome assemblies. Among the seven assemblies, three were generated in the present study (Plhal710_r2, Plhal304_r1, and Plhal703_r1), and four were retrieved from GenBank (available on 12/15/2022). N50 or N75: length of scaffold or contig such that respectively 50% or 75% of the genome is in scaffolds or contigs of this length or longer; L50 and L75: smallest numbers of contigs whose length sums make 50% or 75% of genome size, respectively.

|  | Plhal710_r2 | Plhal304_r1 | Plhal703_r1 | Plhal710_r1 | A23R734 | A23_Pilon | OS-Ph8-99-BIA4 |
| --- | --- | --- | --- | --- | --- | --- | --- |
| **Assembly** | JASSUX000000000 | JASSUW000000000 | JASMMX000000000 | GCA_003724065.1 | GCA_003640465.1 | GCA_004380875.1 | GCA_900000015.1 |
| **Publication** | This work | This work | This work | Pecrix *et al.*, 2018 |  |  | Sharma *et al.*, 2015 |
| **Assembly** | 710r2 | 304 | 703 | 710r1 | A23R734 | A23_Pilon | OS_Ph8_99_BIA4 |
| **Sequencing technology** | PacBio | PacBio | PacBio | Illumina HiSeq | Illumina MiSeq; PacBio | Illumina MiSeq; PacBio | Illumina HiSeq 2000 |
| **Total length** | 74,828,008 | 72,590,531 | 72,058,149 | 67,629,775 | 71,476,700 | 79,878,416 | 74,743,905 |
| **# contigs or scaffolds** | 88 | 122 | 94 | 745 | 7.628 | 1.761 | 3.162 |
| **Largest contig or scaffold** | 4,219,765 | 1,988,880 | 3,552,761 | 1,519,139 | 186.387 | 696.487 | 3,420,761 |
| **GC (%)** | 45.76 | 45.74 | 45.74 | 45.29 | 45.51 | 45.5 | 45.28 |
| **N50** | 1,332,092 | 1,018,182 | 1,220,046 | 4962.78 | 32,620 | 157,860 | 1,546,205 |
| **L50** | 20 | 26 | 19 | 43 | 654 | 154 | 16 |
| **# N's per 100 kbp** | 0 | 0 | 0 | 3,033.21 | 41.58 | 3,143.68 | 11,408.23 |
| **BUSCO** |  |  |  |  |  |  |  |
| **Completeness** | 95.9% | 95.2% | 95.9% | 92.6% | 96.3% | 0.97 | 95.9% |
| **Duplication** | 2.6% | 1.5% | 1.5% | 0.7% | 3.3% | 2.6% | 1.1% |
| **EukCC** |  |  |  |  |  |  |  |
| **Completeness** | 99.16% | 97.48% | 98.32% | 95.8% | 1 | 1 | 1 |
| **Contamination** | 0 | 0.84% | 0 | 0 | 0.84% | 1.68% | 0 |
| **# coding genes** | 16,540 | 16,470 | 16.382 | 36.892 | - | - | 15.459 |

We predicted 1,323 potentially secreted proteins in pathotype 710, and 1,297 in pathotypes 304 and 703, corresponding to about 8% of the proteome for each pathotype (Table S4). Considering proteins with a RxLR and/or WY-domain, we selected 131, 134, and 138 putative effectors in the secretome of Plhal710_r2, Plhal703_r1, and Plhal304_r1, respectively (Tables S5 and S6).

### Structural variation among *Pl. halstedii* pathotypes and synteny with other Peronosporales

Genome alignments show a high level of synteny between all genomes, indicating that the three pathotypes globally share a common genomic backbone (Fig. 1). Pairwise comparisons of the genomes reveal several large-scale inversions ranging from 105 to 932 kb (two between Plhal710_r2 and Plhal703_r1, and four between Plhal710_r2 and Plhal304_r1). We then carried out subsequent analyses using the genome of pathotype 710 (Plhal710_r2) as a reference, as it is the most contiguous.

**Fig. 1.**
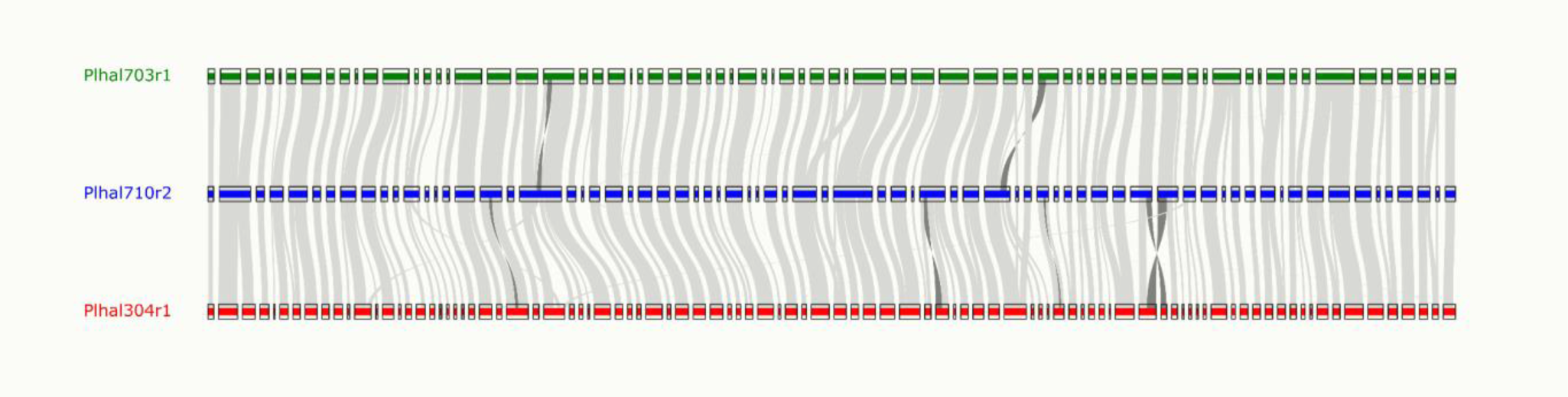
Synteny between the genomes of *Plasmopara halstedii* pathotypes 304, 703, and 710. Genomic blocks were identified and visualised with the MCscan tool implemented in JCVI. Large inversions are highlighted by dark grey links.

The *Pl. halstedii* genome is highly syntenic with the 17 chromosomes of the *Peronospora effusa* telomere-to-telomere assembly (Fig. 2) (Fletcher *et al*., 2022). This points towards a close or identical number of chromosomes between these species, in accordance with the proposed ancestral chromosome number (17) for the Peronosporaceae (Fletcher *et al*., 2022).

**Fig. 2.**
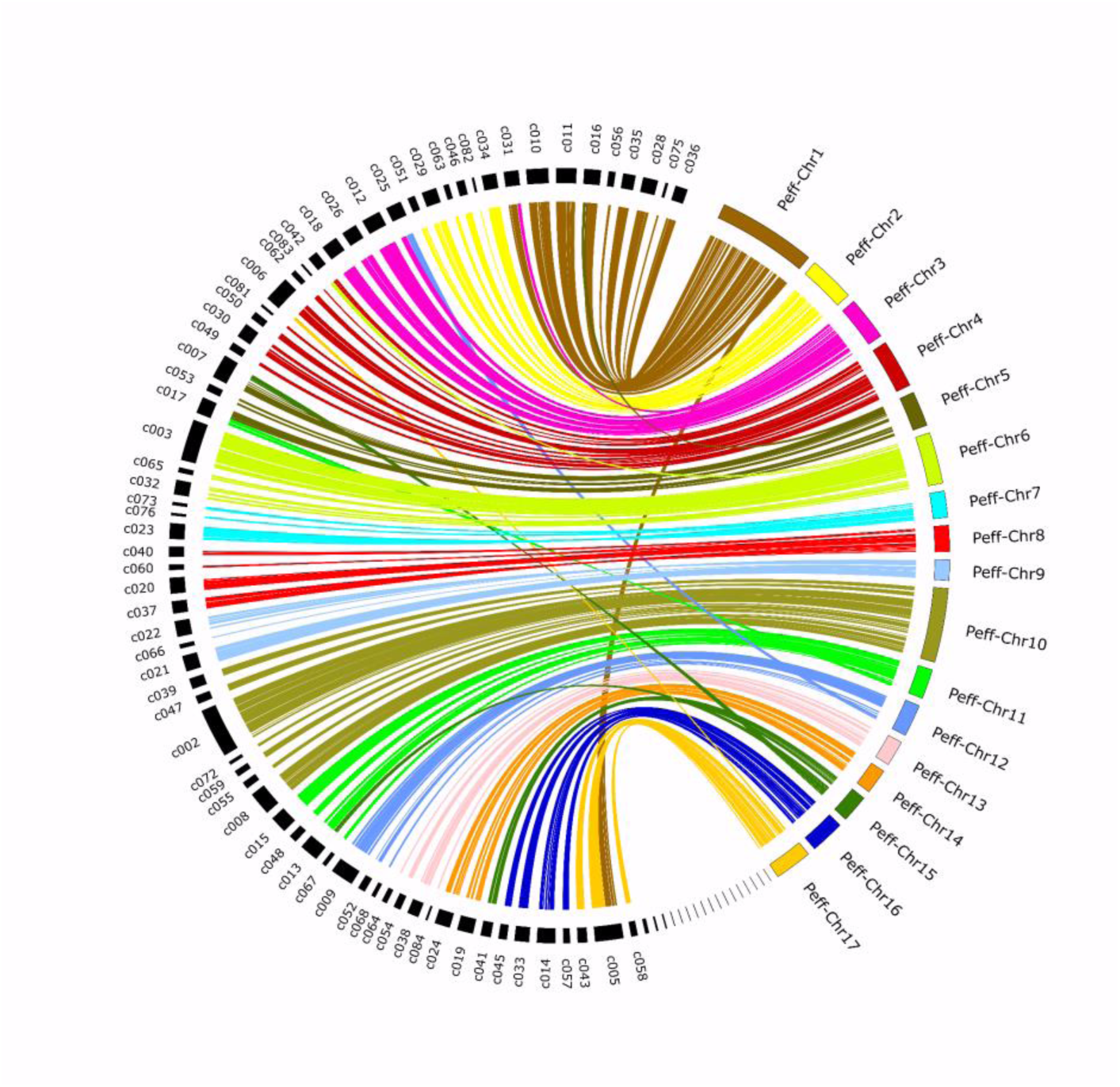
Synteny between the genome of *Plasmopara halstedii* pathotype 710 (Plhal710_r2) with the chromosome-level assembly of *Peronospora effusa* (Fletcher *et al*., 2022). The positions of 5,531 CDS are linked, anchoring 97.5% of the Plhal710_r2 genome length. Links are coloured according to the *Pe. effusa* chromosomes. Corresponding figures for pathotypes 304 and 703 are available in Fig. S2.

### History of the *Pl. halstedii* invasion in France

We performed a population structure analysis based on 97,915 SNPs obtained from next-generation sequencing (NGS) data of the 17 pathotypes (Pecrix *et al*., 2018a) mapped on the Plhal710_r2 reference assembly. STRUCTURE was used to estimate the individual ancestry. The maximum number of clusters obtained was 5, regardless of the higher *K* value (Figs 3a and S3). As a consequence, we divided the 17 *Pl. halstedii* pathotypes in five main genetic clusters. Only seven pathotypes were assigned to a specific cluster (i.e. 710, 100, 300, 304, 703, 700, and 334), while the ten remaining pathotypes were identified as admixed (Fig. 3a). A discriminant analysis of principal components (DAPC) performed on the same SNP dataset showed similar results to those obtained with STRUCTURE (Fig. 3b).

**Fig. 3.**
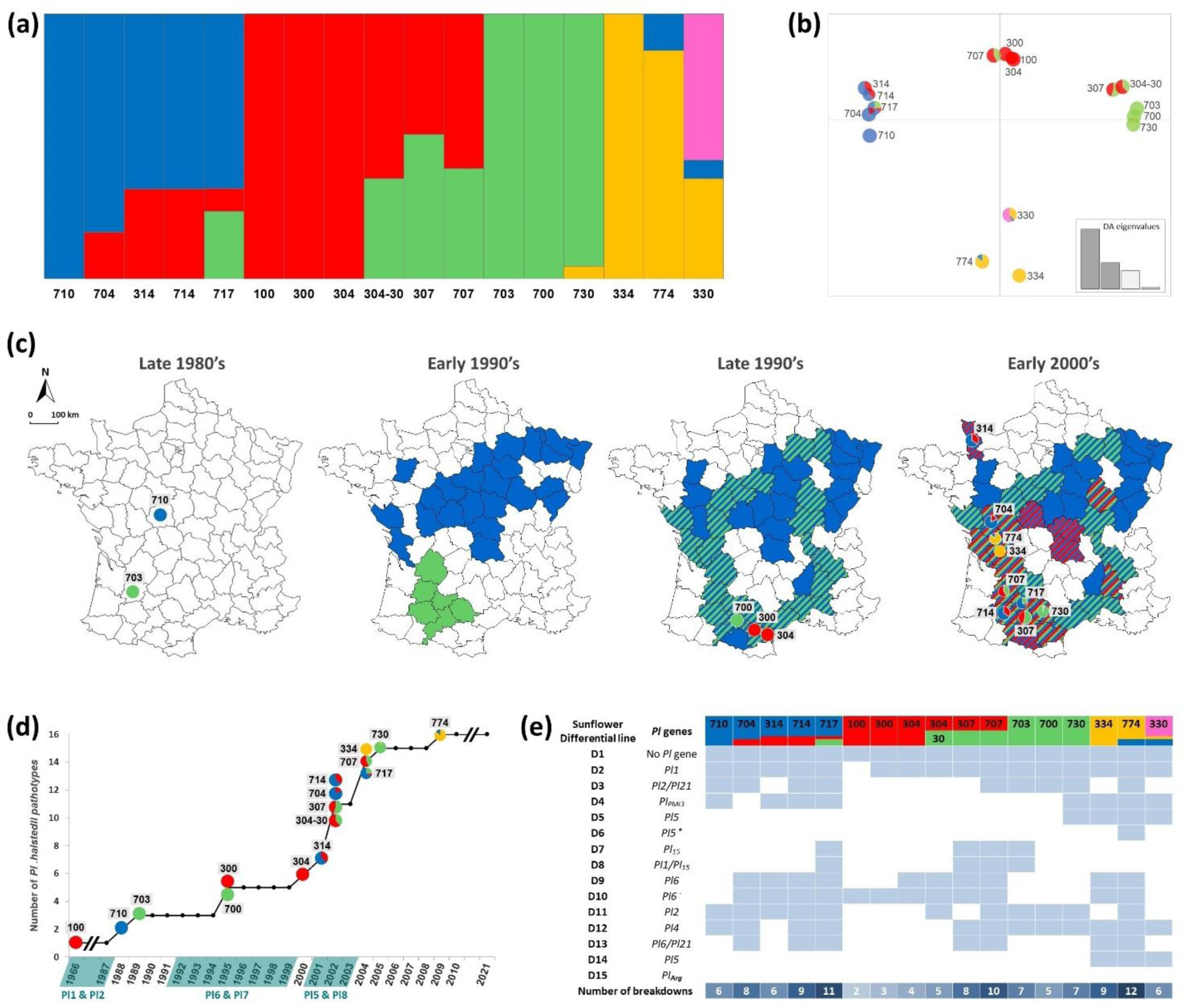
History of the *Plasmopara halstedii* invasion in France: (a) Genetic clustering analysis based on 97,915 SNPs using STRUCTURE for *K* = 5 (see Fig. S4 for other *K* values). Seven pathotypes were assigned to a specific cluster (i.e. 710, 100, 300, 304, 703, 700, and 334), while the ten remaining pathotypes were assigned to two or three clusters, and thus considered as admixed. Colours of genetic clusters here defined their assignment level to each pathotype, are hereafter reused in panels b, c, d, and e. (b) Genomic clustering analysis using a discriminant analysis of principal components (DAPC). Colours in pie charts correspond to the admixture pattern revealed with the STRUCTURE analysis for *K* = 5 (panel a). (c) Spatial distribution of French pathotypes over time (Terres Inovia data; https://www.terresinovia.fr). (d) Emergence of pathotypes and successive breakdowns of sunflower resistances (updated from Ahmed *et al*., 2012). (e) Classification of *Pl. halstedii* pathotypes according to their genetic clusters and virulence profiles (blue squares) on sunflower reference genotypes, adapted from Tourvieille *et al*. (2012) and Gascuel *et al*. (2015).

By integrating genomic structure results with the geographical origins of the pathotypes (Fig. 3c), the timeline of their emergence (Fig. 3d), and their virulence profiles on differential sunflower lines (Fig. 3e), we inferred the invasive history of *Pl. halstedii* in France over the past 70 years. The *Pl. halstedii* pathotype 100 was initially reported in France in 1966 (Moinard *et al*., 2006; Fig. 3d). The first sunflower breeding programs thus introduced two *Pl* resistance genes (i.e. *Pl1* and/or *Pl2*) in all cultivated hybrids sold in France, allowing containment of this pathotype and successful sunflower cultivation for several years. However, two new virulent pathotypes, 710 and 703, were introduced, leading to the breakdown of resistance genes used at the time. These were first described in 1988 and 1989, respectively, in the Centre and South West of France (Moinard *et al*., 2006; Fig. 3c). In the early 90’s, pathotypes 710 and 703 spread widely in France, respectively towards north and south (Fig. 3c), prompting the introduction in sunflower hybrids of new *Pl* genes originating from *Helianthus annuus* (*Pl6*) and *H. praecox* (*Pl7*) (Fig. 3d), both conferring identical range of pathotype resistance. During the late 90’s, three new pathotypes were recorded in southern France: 300, 304, and 700 (Fig. 3c). According to the genomic structure analysis, 300 and 304 are closely related to 100, while 700 is related to 703. Pathotype 304 especially led to the breakdown of *Pl6* and *Pl7* resistance genes. In the same time, 710 and 703 have spread to new areas in France (Fig. 3c) leading to their co-presence in approximately half of the production regions (hatched on Fig. 3c). In the early 2000’s, the prevalence of pathotypes 300 and 304 has also increased in several areas (Fig. 3c,d). The STRUCTURE analysis suggests they have recombined with other pathotypes (Fig. 3a), giving rise to admixed pathotypes 314/704/714 (resulting from the admixture between pathotypes related to 304 and 710), and 304-30/307/707 (resulting from the admixture between pathotypes related to 304 and 703). Pathotype 717 even represents admixture between three genetic clusters (Fig 3a). In total, seven novel recognized pathotypes (all genetically admixed) have thus arisen in the early 2000’s, overcoming *Pl6* and *Pl7* resistance genes. Two other *Pl* genes from LG13, *Pl5* and *Pl8,* were introduced in the meantime in cultivated sunflowers (Fig. 3d; Pecrix *et al*., 2018a) allowing containment of these new pathotypes until 2004 (Fig. 3d,e). In 2004, pathotype 334 was newly recorded, breaking down *Pl5* and *Pl8* sunflower resistance genes (Fig. 3d,e). Pathotype 334 is relatively close to 330 (a Spanish pathotype not yet introduced in France) and constitutes a new genetic cluster (yellow group) on the STRUCTURE analysis. Our analysis suggests that pathotype 334 has then recombined with 703 and 710 relatives giving rise in the late 2000’s to pathotypes 730 and 774 (Fig. 3).

Thus, in line with the results of population genomics and the chronology of the *Pl. halstedii* emergence in France, we considered pathotypes 100, 334, 703, and 710 as representatives of the founder lineages at the origin of all the pathotypes showing admixture signals. Since pathotype 100 is no longer present in France, we selected the closely related pathotype 304 as the founder representative of the red group.

### Genetic admixture of *Pl. halstedii* pathotypes and the emergence of new virulences

Population structure indicates that admixture events have played a decisive role in the emergence and establishment of numerous virulent *Pl. halstedii* pathotypes in France, and have enabled their adaptation to cultivated hosts. By combining the phenotypic traits of native or introduced parental strains, hybridization events have generated new pathotypes capable of infecting a greater number of resistant sunflower lines (Fig. 3e). In the majority of cases, admixed pathotypes have a wider range of virulence among differential lines than the parental pathotypes, suggesting that admixture events had a positive effect on the ability to overcome resistance (Fig. 3e). Pathotypes 704, 314, and 714 are able to infect eight, six, and nine differential lines, respectively, whereas parental pathotypes 304 and 710 are virulent on only four and six. Similarly, pathotypes 304-30, 307, and 707 are able to infect five, eight, and ten lines, respectively, compared to four and seven for parental pathotypes 304 and 703. The most striking case concerns pathotype 717, resulting from the admixture of the three parental strains 304, 703, and 710, which is able to infect 11 of the differential lines. Interestingly, these hybridizations did not result in an increased heterozygosity in admixed strains. The estimations of homozygosity rate are similar between founder and admixed strains, ranging from 60.1% to 87.8% of SNPs (Fig. S5).

To explore the genetic exchanges that structured the genomes of *Pl. halstedii* strains, we generated an ancestral recombination map for each admixed pathotype (Fig. 4). The method consisted of reconstructing the ML phylogeny of each 100 kb non-overlapping window across the *Pl. halstedii* genome. We thus partitioned the genome into 100 kb non-overlapping windows, built a ML phylogenetic tree for each genomic window, and identified the closest relative of each admixed pathotype (according to STRUCTURE) among the four genetically non-admixed strains (i.e. 100, 334, 703, and 710). We obtained 811 phylogenies which we classified into five groups based on the phylogenetic proximity of the admixed strain with its closest non-admixed relative (Fig. 4a and Fig. S3). Consistently with STRUCTURE, the admixed pathotypes 300 and 304 clustered with the parental pathotypes 100 across the whole-genome (red topologies). Considering the admixed pathotypes 314, 704, and 714 resulting from hybridisation between parental strains 710 and 100, we consistently observed two topologies, i.e. clustering either these pathotypes with strain 710 (blue topologies) or with strain 100 (red topologies). Similarly, the admixed pathotypes 304-30, 307, and 707 are clustered in the majority of cases with one of their inferred parents, i.e. either pathotype 100 (red topologies) or 703 (green topologies). This phylogenomic approach confirmed the inferred population structure, highlighted the genomic mosaicism of admixed pathotypes, and revealed the genomic regions they share with their founder pathotypes. Deployment of new resistances thus seems to be correlated with wide chromosomal exchanges in admixed pathotypes that occurred in the past (Fig. 4).

**Fig. 4.**
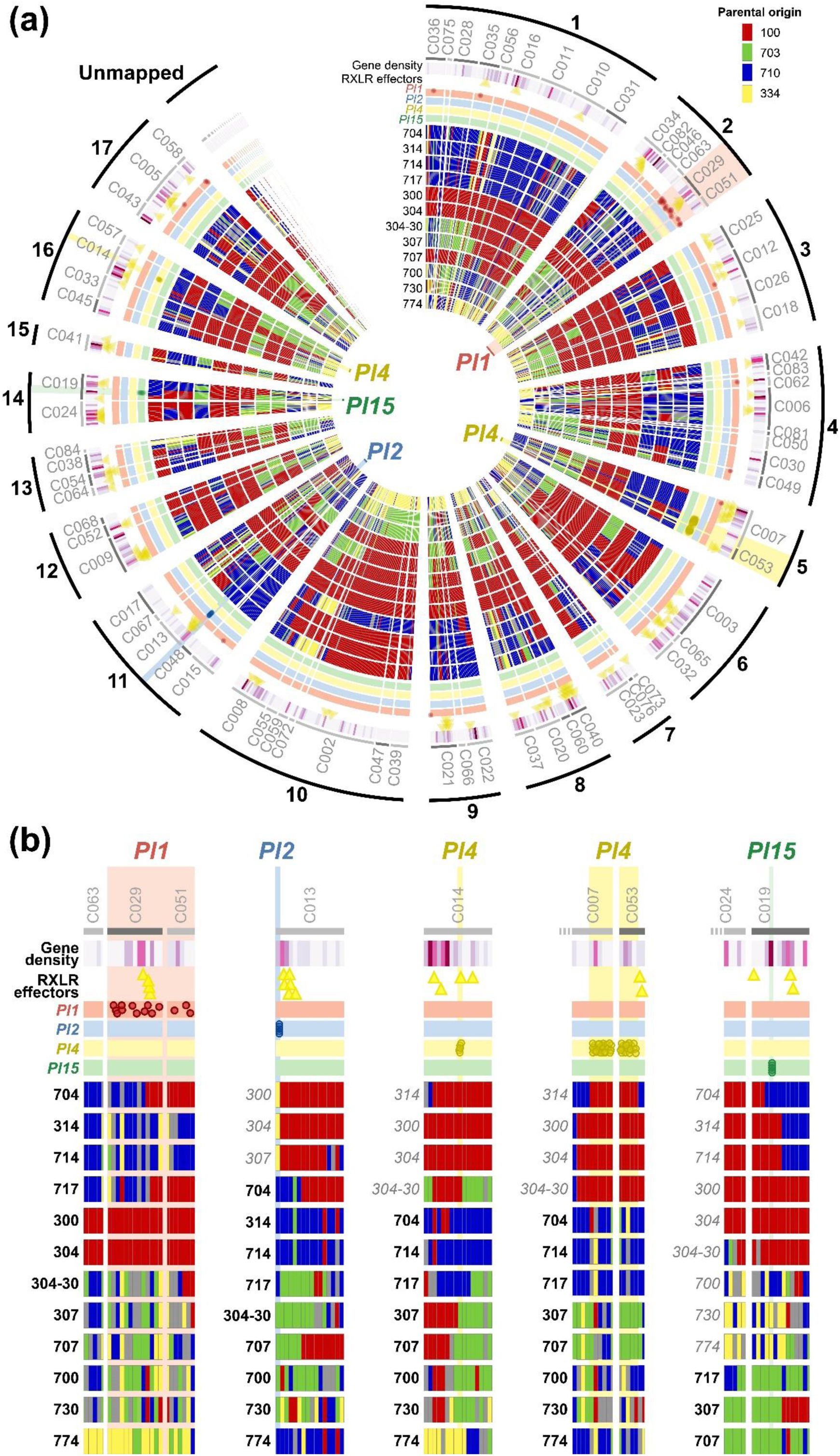
Genome wide integrative view of the *Plasmopara halstedii* Plhal710_r2 reference genome with phylogenomic data of French pathotypes and genomic regions associated with sunflower resistance gene breakdowns. The red, green, blue, and yellow trees correspond to the topologies for which an admixed strain is the closest relative to the parental pathotype 100, 703, 710, or 334, respectively (Fig. S3). (a) Circular representation of the *Pl. halstedii* Plhal710_r2 reference genome. From the outside to the inside of the Circos plot: the external ring represents Plhal710_r2 contigs gathered into pseudo-chromosomes inferred via synteny with *Pe. effusa* chromosomes; The second ring shows the contigs of Plhal710_r2 assembly ordered and oriented to constitute pseudo-chromosomes. The contigs that have been inverted to respect the synteny with *Pe. effusa* are coloured dark grey. The heatmap rings represent the density of genes coding secreted proteins. In the following ring, RxLR effectors are shown as yellow triangles. The red, blue, yellow, and green rings show the distribution of SNPs associated with the breakdown of *Pl1*, *Pl2*, *Pl4*, and *Pl15*, respectively. The 12 inner rings are the chromosome paintings of admixed pathotypes representing the topologies for each 100-kb non-overlapping window across the chromosomes. (b) Detailed view of genomic regions associated with resistance breakdown. For each region, the virulent pathotypes are indicated in black, and the avirulent ones in grey italics.

To investigate the genetic determinisms of *Pl* resistance gene breakdowns, we explored the mosaic regions of admixed pathotypes. For each of these resistances, we first looked for polymorphic variation between virulent and avirulent pathotypes. With this approach, we identified 25, 12, 171, and five SNPs associated with the breakdown of *Pl1*, *Pl2*, *Pl4*, and *Pl15*, respectively (Fig. 4). Strikingly, most of these SNPs are concentrated in a few genomic regions of 1.47 Mb, 12.6 kb, 0.86 Mb, and 2.95 kb for the *Pl1*, *Pl2*, *Pl4*, and *Pl15* breakdown (Fig. 4b), respectively. Interestingly, these SNPs overlap with mosaic regions of the admixed pathotypes (Fig. 4b, Table S7). These regions are of particular interest, as they may contain genes associated with resistance breakdown and will be the main focus of our subsequent analyses.

All SNPs associated with *Pl2* resistance are distributed across one genomic window where avirulent pathotypes (300, 304, and 307) cluster with the avirulent founder strain 334 (yellow topologies) and where virulent pathotypes cluster with the virulent founder strains 710 (blue topologies) or 703 (green topologies). Similarly, all SNPs associated with *Pl4* resistance are distributed across genomic windows where avirulent pathotypes (300, 304, 304-30, and 314) cluster with the avirulent founder strain 100 (red topologies) and where virulent pathotypes cluster with the virulent founder strains 710 (blue topologies) or 703 (green topologies). Consistently, all SNPs associated with *Pl5* resistance are distributed across one genomic region where virulent pathotypes (307, 707, and 717) cluster with the virulent founder strain 703 (green topologies) and where avirulent pathotypes cluster with the three other founder strains. In contrast, regions associated to *Pl1* breakdown could not be investigated further since no high-quality assembly of pathotype 100, the only avirulent pathotype on *Pl1*, is available, and therefore no genomic comparison could be made with virulent pathotypes.

### Towards identification of candidate genes responsible for breakdown of resistance genes

In this section, we attempted to identify the genetic determinants driving the emergence of pathotypes with new virulence patterns. We first identified genes encoding putative secreted effectors containing the RxLR motif and/or WY domains in the three founder genomes by *in silico* classical methods: 131, 138, and 134 putative effectors were found in Plhal710_r2, Plhal304_r1, and Plhal703_r1 genomes, respectively (Tables S4 and S6). The 131 putative effectors from pathotype 710 were localised on the genomic contigs (Fig. 4). Most of the predicted effector proteins are strongly conserved in the other two pathotypes, 73% and 51% of 710’s effectors being completely identical in 703 and 304 pathotypes, respectively (Table S6). However, three putative effectors of 710 show less than 40% of identity in 703, which is not the case with 304 (98% identity), suggesting a bigger distance between pathotype effectors of 710 and 703 than with 304, as shown on the DAPC for the complete genomes (Fig. 3). Based on SNPs, we mapped the regions of interest (ROI) associated with resistance breakdown (Fig. 4b) on the contigs of the three founder pathotypes and listed all the coding genes. Among them, we focused on sequences encoding secreted proteins and putative effectors.

In the ROI of *Pl2* resistance breakdown, the virulent 710 and 703 pathotypes have only non-coding (nc) RNAs (32 for 710, and 12 for 703), whereas avirulent 304 has in addition one complete coding gene, Plhal304r1c045g0126491 (Table S7). The 112 amino acids encoded protein of this gene is not predicted to be secreted or to be an effector and is present in both virulent 710 and 703 pathotypes, on the border of two other contigs. Therefore, candidate genes associated with *Pl2* breakdown were not identified. However, the ROI corresponds to the extremity of a contig in the Plhal710_r2 and Plhal304_r1 assemblies, raising the possibility that a part of the region is missing or incorrectly assembled.

In the ROI for *Pl4* breakdown, the avirulent pathotype 304 has 231 encoded proteins, while the virulent 710 and 703 pathotypes have 234 and 227 proteins, respectively (Table S7). Twenty-one (304, 710) and 23 (703) proteins are predicted to be secreted in the ROI. None of them display the RxLR motif but one, Plhal304r1c027g0090151, has five WY domains suggesting it could be indeed an effector (Table S7). Alignment with the homologous proteins of pathotypes 710 and 703 shows two substitutions and the gain of 17 amino acids in the middle of the sequence (Fig. S6). Therefore, this protein is a promising candidate effector involved in the interaction with the *Pl4* resistance gene.

In the ROI for *Pl15* breakdown, the avirulent pathotypes 304 and 710 have 150 and 181 protein coding genes, respectively, and among them 16 and 22 secreted proteins. The 703 virulent pathotype has 181 proteins in the ROI region and 23 secreted proteins (Table S6). The only RxLR gene in the *Pl15* ROI that was predicted *in silico* with the method of Bhattacharjee *et al*. (2006) is Plhal710r2c019g0083331. The gene is, however, 100% conserved in the virulent 703 pathotype (Plhal703r1c26g0109591) and can thus not explain the *Pl15* breakdown. Another candidate is Plhal710r2c019g0082461, which was already annotated by Sharma *et al*. (2015) as PHALS_04751. This protein is identical in pathotype 304 (Plhal304r1c010g0039761). Interestingly, in the syntenic region of pathotype 703, we found a sequence coding for a protein of similar length but very divergent (32.7% identity; Fig. S7). The 703 protein is predicted to be secreted, which is not the case for the allelic forms of 710 and 304. However, if we consider that the protein starts were mis-predicted and started at the 23rd amino acid, a signal peptide can be predicted (aa23 to aa44; Fig. S7). In our pathotypes panel, all virulent strains possess the 703 gene, while avirulent genotypes present the 710/304 gene with the exception of pathotype 700 that appears heterozygous at the locus. This gene is therefore a good candidate for a gain of virulence on the *Pl15* resistance gene.

## DISCUSSION

Our study compares the genetic structure, the genomic organization, and the virulence profile among 17 *Pl. halstedii* pathotypes introduced in western Europe (including the 16 reference ones described in France). By integrating these data, we reconstructed the history of the pathogen in France over the last 60 years, and investigated how emerging pathotypes were able to break down sunflower resistance genes. The population genomic analyses demonstrated that, from the introduction of a few strains with contrasting genetic backgrounds, the pathogen has mainly evolved by admixture, resulting in new virulence profiles. To investigate the role of recombination in the evolution of virulence, we performed a phylogenomic analysis of the whole-genomes. The analysis of the genomic mosaicism of admixed pathotypes identified candidate avirulence regions associated with resistance breakdown. Altogether, these results provide a better understanding of the genetic bases underlying the adaptation of the pathogen to its host plant.

### High-quality genome assemblies reveal conserved synteny among Peronosporales and low heterozygosity in *Pl. halstedii*

The first step of our study was the production of high-quality genomes of the sunflower downy mildew. We thus provide three long reads genomes, which are more complete and better resolved than previous genomes generated with short reads only (Sharma *et al*., 2015; Pecrix *et al*., 2019). The high level of contiguity of our assemblies allows the comparison to other genomes of oomycete pathogens, in particular with those of downy mildews of grapevine (*Pl. viticola*) and spinach (*Peronospora effusa*). A strong synteny with other Peronosporales species was observed, confirming a genome organization in 17 chromosomes (Fletcher *et al*., 2022; Dussert *et al*., 2019; Dvorak *et al*., 2025b). It is important to point out that, in numerous cases, SNPs associated with resistance breakdowns are concentrated on only one pseudo-chromosome, which reinforces the reliability of the pseudo-chromosomes inferred from synteny with *Pe. effusa*.

A low heterozygosity was observed in all *Pl. halstedii* genomes confirming that this oomycete species should mainly self-fertilize during its sexual reproduction phases (Spring, 2000; Ahmed *et al*., 2012; Gascuel *et al*., 2015). Indeed, heterozygosity estimates range from 120 (Sharma *et al*., 2015) to 141-166 heterozygous sites/Mb (in this study), which are very far from those observed in the strictly heterothallic *Pl. viticola* (mean of 7,952 heterozygous sites/Mb; Dussert *et al*., 2019). Heterothallism (i.e. sexual reproduction requiring individuals with different mating types) leads to out-crossing in *Pl. viticola* (Wong *et al*., 2001), while in *Pl. halstedii*, homothallism should largely favour self-fertilization. However, this does not exclude occasional out-crossing between strains, a mechanism that allows the reshuffling of allelic combinations. After such admixture events, new genomic combinations in *Pl. halstedii* are probably rapidly fixed by self-fertilization and/or inbreeding between very closely related strains.

### Monitoring and evolution of virulence in *Pl. halstedii* populations

Invasive plant pathogens in agro-ecosystems provide useful models for understanding the evolutionary processes that lead to the emergence of new virulence. Sunflower downy mildew is the subject of particular attention since the 60’s from the French plant protection services, which have been called upon to monitor the progression of the pathogen in France. There are very few examples of such historical monitoring of a plant pathogen emergence, documented over a long period of time and on a wide geographical scale.

The integrated results from ground monitoring, phenotyping and genotyping of *Pl. halstedii* isolates in France support the hypothesis that at least four successive introductions of the pathogen occurred in France (Louvet & Kermoal, 1966; Delmotte *et al*., 2008; Ahmed *et al*., 2012; Gascuel *et al*., 2015; ANSES, 2016). The *Pl. halstedii* pathotype 100 was initially reported in France in 1966 (Moinard *et al*., 2006). This situation remained stable until the 80s, when two new virulent pathotypes (710 and 703) were introduced in quick succession in 1988 and 1989, respectively, in the Centre and South West of France (Moinard *et al*., 2006). In 2004, a new pathotype 334, close to the Spanish race 330, was reported in region Nouvelle-Aquitaine suggesting a fourth introduction in France.

Previous studies have shown that *Pl. halstedii* pathotypes issued from different introduction events can locally co-exist (Elameen *et al*., 2022; Kitner *et al*., 2023), which should offer the possibility of genomic recombination between them. Our results confirm this hypothesis of admixture in relation to the emergence of new and endemic pathotypes in response to host resistance (Ahmed *et al*., 2012). Between 1987 and 2008, the monitoring revealed a rapid increase in the number of pathotypes rising from one to 14, suggesting an acceleration of pathotype emergence. A large part of the newly emerging pathotypes (e.g. 304, 307, 704, and 714), which have never been recorded outside France, had indeed less than 70% of ancestry assigned to one or other genetic clusters. These pathotypes were all able to overcome *Pl6* resistance of the host. Although we were able to demonstrate how new virulent pathotypes have emerged, we were not able to ascertain from these data how frequently such events occur, or whether crosses between some pathotypes are more likely than others.

### Evidence for virulence evolution by mutation and admixture

New endemic pathotypes presenting novel virulence profiles could arise through mutation occurring in a clonal lineage, a ubiquitous process that has been described in many plant pathogens (e.g. McDermott & McDonald, 1993). Here, the genomic data provides evidence that, in some cases, virulence emergence should have occurred through mutations in a unique *Pl. halstedii* lineage. In the genetic cluster comprising pathotypes 100, 300, and 304, the breakdown of the *Pl1* gene by strains 300 and 304 could be the result of mutations in pathotype 100, which was initially present in France in the 60s. Within the genetic group initially composed of pathotypes 700 and 703, mutations may have enabled it to overcome the *Pl5* gene, leading to the emergence of pathotype 730 in the 2000s.

Despite being rare, out-crossing may be a key component of the evolution of resistance-breaking genotypes as shown here by the emergence of new virulent admixed pathotypes. Strains of *Pl. halstedii* display large structural variations in their genome. Such intra-chromosomal structural rearrangements have also been described in *Ph. infestans* (van der Lee *et al*., 2004; Matson *et al*., 2022) and *Pe. effusa* (Skiadas *et al*., 2024). This structural diversity could contribute to the adaptability of the pathogen population to disease management methods. Interestingly, additional analyses of shotgun sequencing data revealed signatures of widespread recombination events between the three founder pathotypes among a set of isolates representative of the French sunflower downy mildew diversity. Investigating shared genomic regions among admixed pathotypes offer the possibility to apply admixture mapping to identify genomic regions associated to traits (e.g. Buerkle & Lexer, 2008; Fetter & Keller, 2023), and this should bring new insights on *Pl. halstedii* adaptation especially regarding the deployment of their new virulence genes.

### Identification of candidate genes for resistance breakdown

Several strategies can be considered to identify genes associated with virulence and resistance breakdown. Genome-wide association studies (GWAS) has been already successfully applied in some oomycete species to study various life history traits (Dussert *et al*., 2020; Vogel *et al*., 2021; Paineau *et al*., 2024), including virulence towards plant resistance genes. However, the strong genetic structure among *Pl. halstedii* pathotypes coupled with selfing is expected to reduce the power of GWAS due to confounding effects from ancestry differences. QTL (Quantitative Trait Loci) mapping could therefore be explored using experimental crosses to identify genomic regions associated with virulence in *Pl. halstedii*. It has indeed proven effective for other oomycetes, such as *Hyaloperonospora arabidopsidis* (Asai *et al*., 2018), *Ph. sojae* (Arsenault-Labrecque *et al*., 2022), and *Pl. viticola* (Dvorak *et al*., 2025a). However, crossing is technically challenging in obligate biotrophic oomycetes such as *Pl. halstedii*, which cannot be cultured outside their hosts.

In this study, based on SNP analysis and the history of *Pl. halstedii* pathotypes in France, we identified two candidate effector genes breaking the *Pl4* and *Pl15* downy mildew resistances. Regarding the *Pl4* breakdown, the candidate gene encodes a protein with a 17 amino-acid insertion in the virulent pathotypes which could prevent the recognition of this putative effector by *Pl4*. In the case of *Pl15* breakdown, most SNPs linked to admixture from the virulent pathotype 703 are concentrated in one gene coding for a small protein predicted to be secreted. The comparison of the orthologous sequences revealed two highly divergent allelic sequences. All virulent strains possess the 703 allele, while avirulent pathotypes share the divergent form, with the exception of avirulent pathotype 700 that appears heterozygous, suggesting that one avirulent allele is sufficient to trigger recognition and resistance to that strain.

### Concluding remarks and perspectives

Our study shows that whole-genome data can improve our understanding of the evolutionary epidemiology of plant pathogens, particularly by helping to identify admixture events. The identification of several genomic regions associated with resistance breakdown highlights the importance of studying multiple assemblies to adequately study the diversity of plant pathogens (Chen *et al*., 2023; Skiadas *et al*., 2024).

Genome-wide local ancestry approaches offer a powerful framework for understanding the genetic diversity of pathogens, particularly in identifying virulence genes that drive disease emergence and adaptation (Tichkule *et al*., 2021; Guo *et al*., 2022). By characterizing the genomic regions inherited from different parental backgrounds, variations associated with pathogen virulence and host adaptation can be identified. This knowledge enables the targeted and reasoned deployment of resistance genes adapted to the genetic diversity of pathogen populations, thereby reducing the risk of resistance breakdown. Furthermore, by predicting the evolutionary trajectories of pathogen populations, these approaches provide valuable insights into the durability of resistance strategies. Ultimately, integrating genome-wide local ancestry data into disease management opens new perspectives for sustainable agricultural practices, fostering the development of long-lasting resistances and more effective disease control.

## Data availability

The three genomes were deposited at DDBJ/ENA/GenBank under accessions JASSUX000000000 (race 710), JASSUW000000000 (race 703), and JASMMX000000000 (race 304), respectively. The corresponding PacBio RSII raw reads are available in the SRA database under studies SRP442823 (race 710), SRP442826 (race 703), and SRP442827 (race 304). The Illumina resequencing reads were deposited in the SRA database under study SRP443016. The VCF file generated and used in this study is available at data.gouv.fr: https://doi.org/10.57745/XQUTTT.

## Acknowledgments

The authors thank Patrick Vincourt for the Illumina genomic sequencing of *Pl. halstedii* pathotypes and GetPlage platform facilities for excellent paid sequencing services. They thank Emmanuelle Mestries from Cetiom (Terres Inovia) and the French public organism “Protection des Végétaux” for organising the annual surveys and writing reports on *Pl. halstedii* pathotypes in France. The authors are grateful for former research teams led by Denis Tourvieille and Felicity Vear, at INRAE Clermont-Ferrand, who initiated and conducted pathological studies on sunflower downy mildew in France.

This work was supported by the Inter-Units grant EVOPLASMO funded by the French Laboratory of Excellence project “TULIP” (ANR-10-LABX-41; ANR-11-IDEX-0002-02) and the Agrobiosciences, Interactions and Biodiversity Research Federation (FR AIB, FR3450). GB is a member of the CRBE laboratory supported by the French Laboratory of Excellence (LabEx) projects CEBA (ANR-10-LABX-25-01) and TULIP (ANR-10-LABX-41), managed by the French ANR. YP and FL were also supported by the European Regional Development Fund and Région Réunion (ERDF fund 2024-1248-005756). YP and FL acknowledge the Plant Protection Platform (3P, IBiSA).

## Author contributions

YP, GB and LG designed and performed the research; YP, ED, FL, LL, SC, JG, FD, GB and LG performed the analyses; YP, ED, FL, FD, GB and LG wrote the manuscript; all authors edited the manuscript.

## Notes

https://doi.org/10.57745/XQUTTT

https://github.com/fredericlabbe/Phalstedii_Phylogenomics

### Competing Interest Statement

The authors have declared no competing interest.

### Summary of Updates

This version of the manuscript has been revised to update the data accession numbers.

