## Supplementary material for "Phylogenomics of *Plasmopara halstedii* reveals genomic regions associated with the breakdown of sunflower downy mildew resistance genes": Supplementary_Information.docx

### *New Phytologist* Supporting Information

Article acceptance date: Click here to enter a date.

The following Supporting Information is available for this article:

**Fig. S1** Examples of structural variations among *Plasmopara halstedii* sequenced pathotypes.

**Fig. S2** Synteny between the genomes of *Plasmopara halstedii* pathotypes with the chromosome-level assembly of *Peronospora effusa* (Fletcher et al., 2022).

**Fig. S3** The twelve topologies used to investigate the genome-wide local ancestry among French pathotypes.

**Fig. S4** Population genetic structure of *Pl. halstedii* populations in France.

**Fig. S5** Heterozygosity rates of French *Pl. halstedii* pathotypes.

**Fig. S6** Sequence alignment of a candidate effector protein associated with *Pl4* breakdown.

**Fig. S7** Sequence alignment of a candidate effector protein associated with *Pl15* breakdown.

**Table S1** Assembly process summary

**Table S2** Multi-criteria contamination detection

**Table S3** Illumina datasets

**Table S4** *Plasmopara* *halstedii* effector statistics

**Table S5** Secretomes

**Table S6** Effectomes

**Table S7** Summary of regions of interest for resistance breakdowns

[Note: if your file is a large table, e.g. an Excel file, this should be submitted separately]

**Supplemental Figures**

**Fig. S1 Examples of structural variations among *Plasmopara halstedii* pathotypes. (a)** 280 kb-long inversion between contigs Plhal710r2c008 (Phal710_r2) and Plhal304r1c003 (Phal304_r1). (**b)** 520 kb-long inversion between contigs Plhal710r2c002 (Phal710_r2) and Plhal703r1c002 (Phal703_r1). Large inversions of gene order between pathotypes are indicated in red. Alignment generated with D-GENIES (Cabanettes & Klopp, 2018).

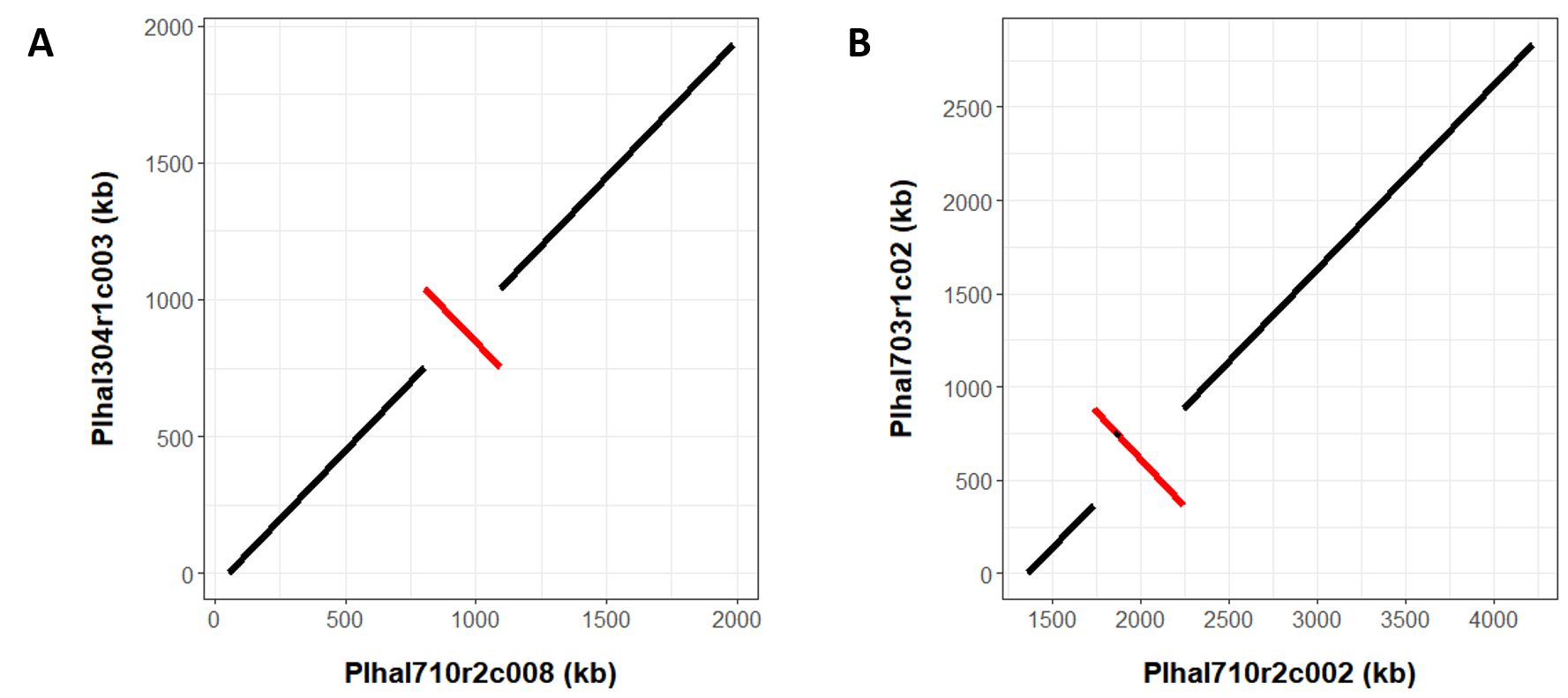

**Fig. S2** Synteny between the genomes of *Plasmopara halstedii* pathotype Phal304_r1 (a) and Plhal703-r1 (b) with the chromosome-level assembly of *Peronospora effusa* (Fletcher *et al.*, 2022). Links are colored according to the *P. effusa* chromosomes on the right of the panels.

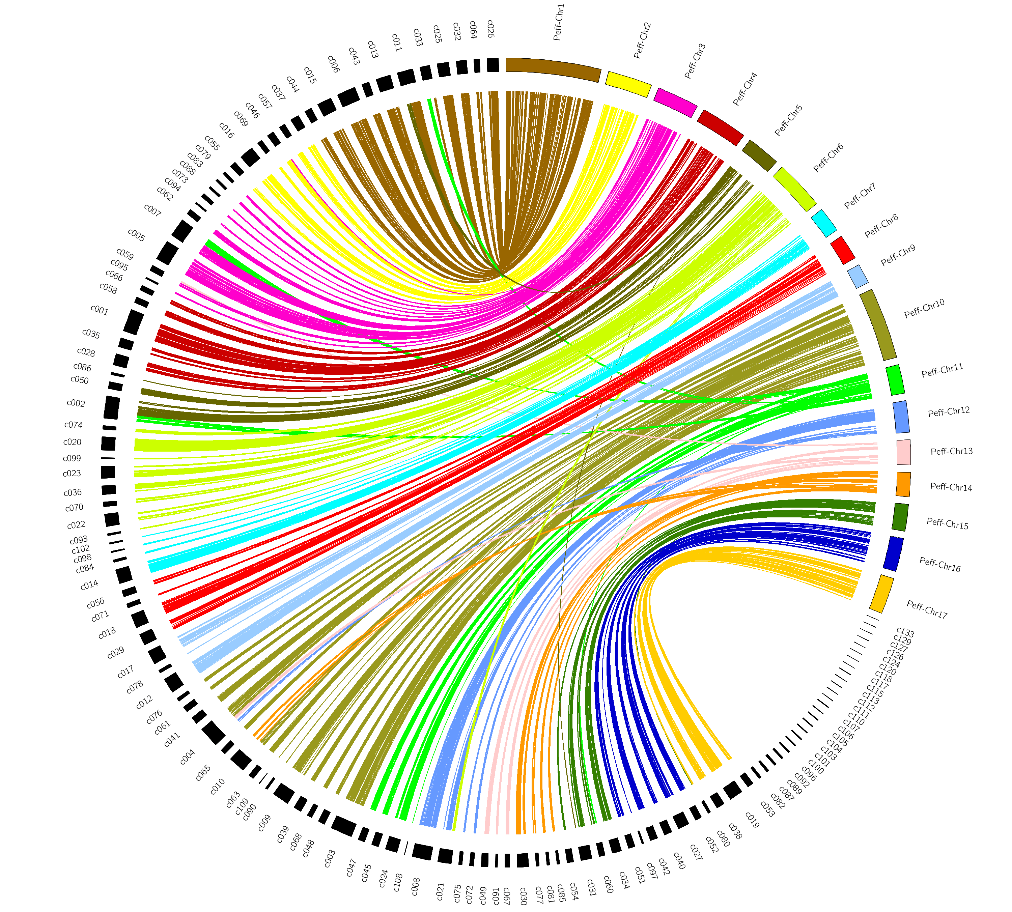

**(a)**

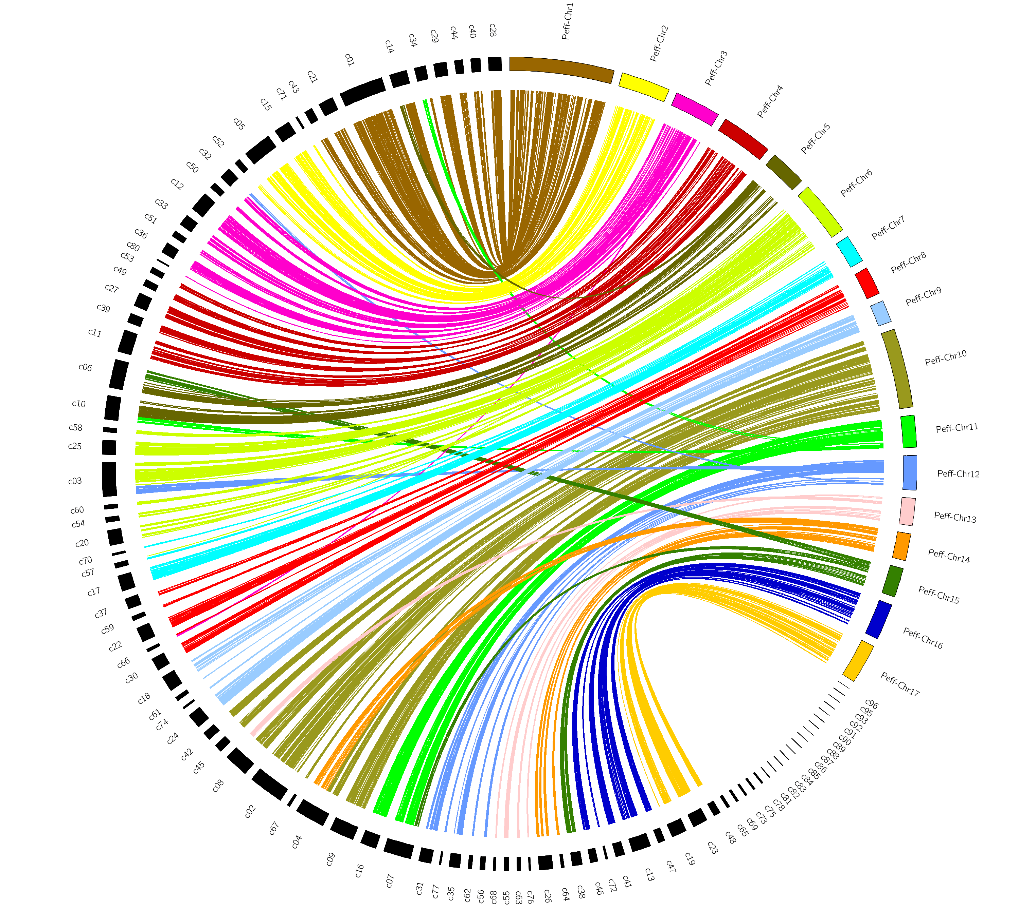

**(b)**

**Fig. S3 The twelve topologies used to investigate the genome-wide local ancestry among French pathotypes.** Phylogenetic trees were reconstructed in 100 kb non-overlapping windows along the chromosomes using PhyML. The red, green, blue, and yellow topologies correspond to those for which an admixed pathotype (*i.e.,* the tree branch without a leaf) is the closest relative to the parental pathotypes 100 (PLHAL100), 703 (PLHAL703), 710 (PLHAL710), or 334 (PLHAL334), respectively. The three topologies in which an admixed pathotype has no closest relative among the parental pathotypes are not shown.

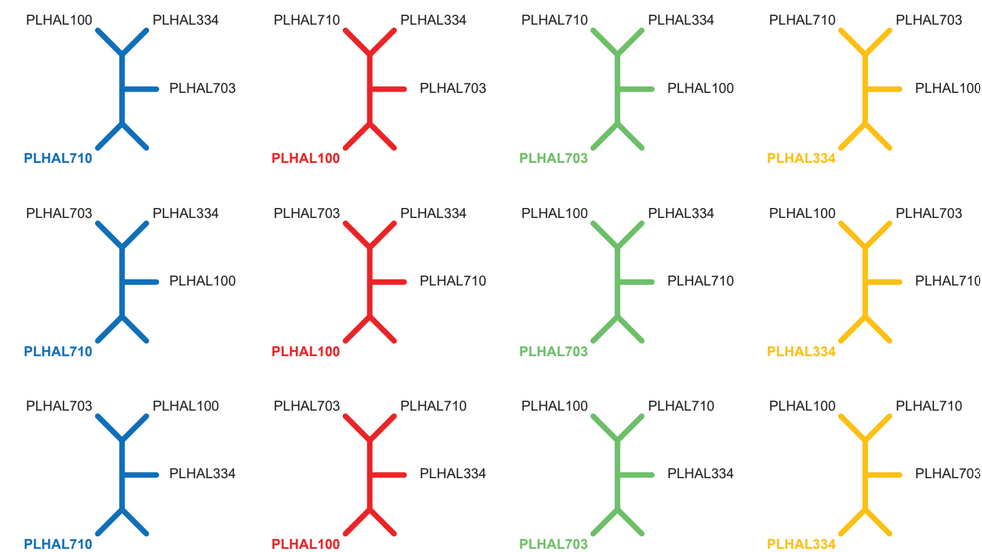

**Fig. S4** Population genetic structure of *Pl. halstedii* populations in France. In this analysis performed using STRUCTURE (Pritchard et al., 2000), each pathotype is represented by a vertical bar segmented with different colored genetic clusters, with length proportional to the probability of assignment to each cluster. The analysis, which tested cluster numbers from K=3 to K=10, identified K=5 as the most likely number of clusters, as all pathotypes were consistently divided into 5 clusters regardless of higher K values.

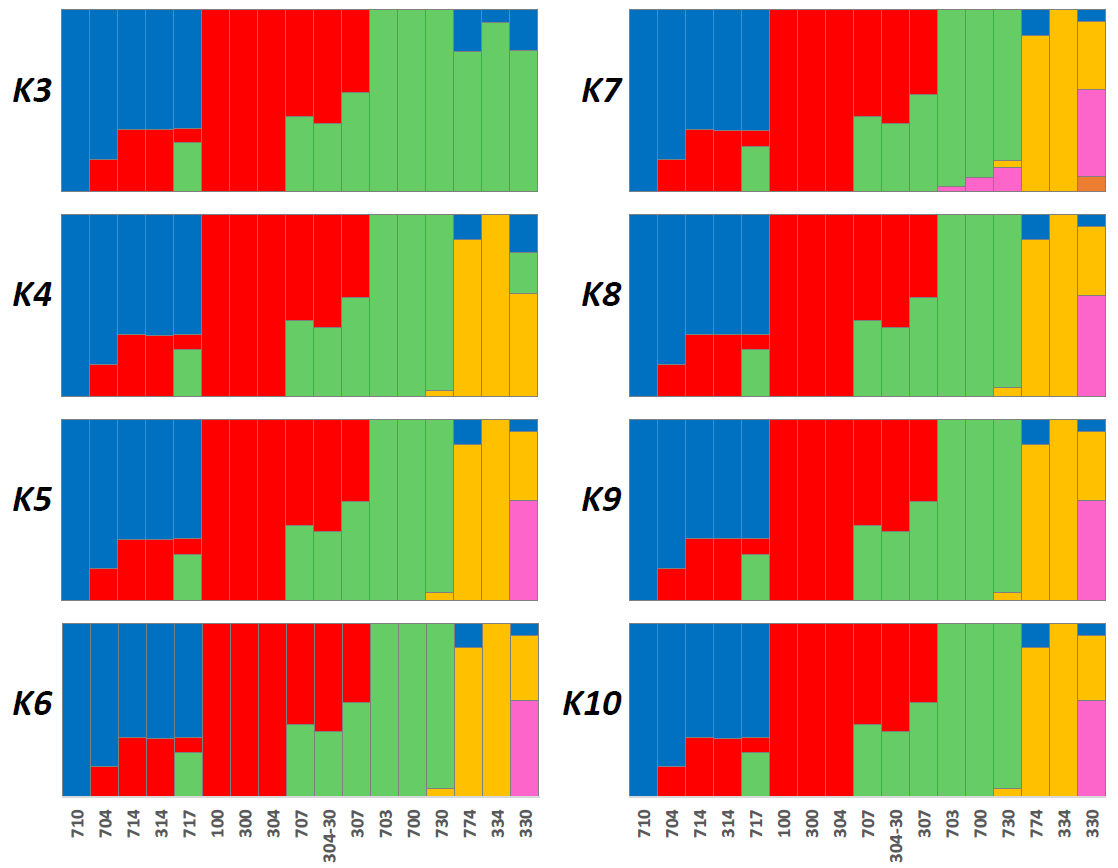

**Fig. S5** Heterozygosity rates of French *Pl. halstedii* pathotypes. Estimates of heterozygosity rates are based on 97,915 bi-allelic SNPs that can be categorized as homozygous (convergent or divergent from reference 710), heterozygous or missing.

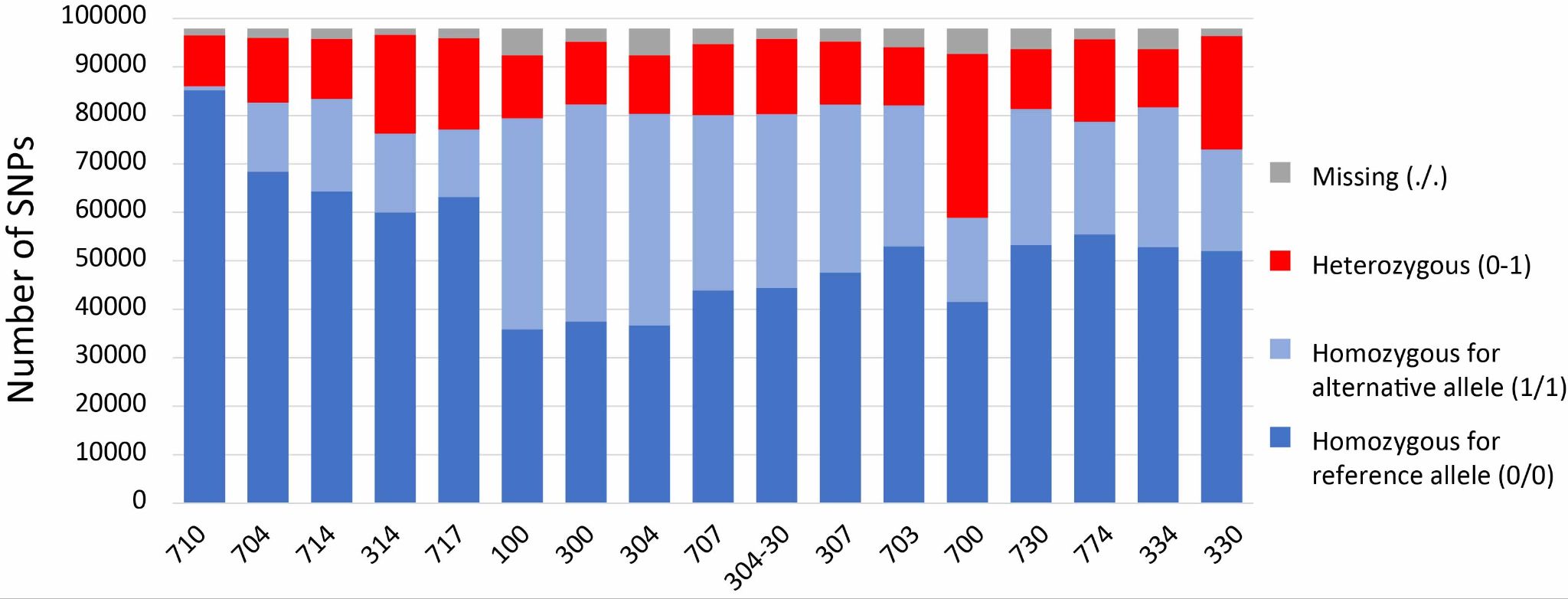

**Fig. S6** Sequence alignment of a candidate effector protein associated with *Pl4* breakdown. The Pl4-avirulent allele is from 304 pathotype (Plhal304r1c027g0090151 gene), whereas Pl4-virulent allele from pathotypes 703 (Plhal703r1c13g0066451 gene) and 710 (Plhal710r2c014g0063611 gene) carries an identical 17 amino acid insertion and 2 amino acid changes compared to Pl4-304 avirulent allele. Divergent amino acids are highlighted in color.

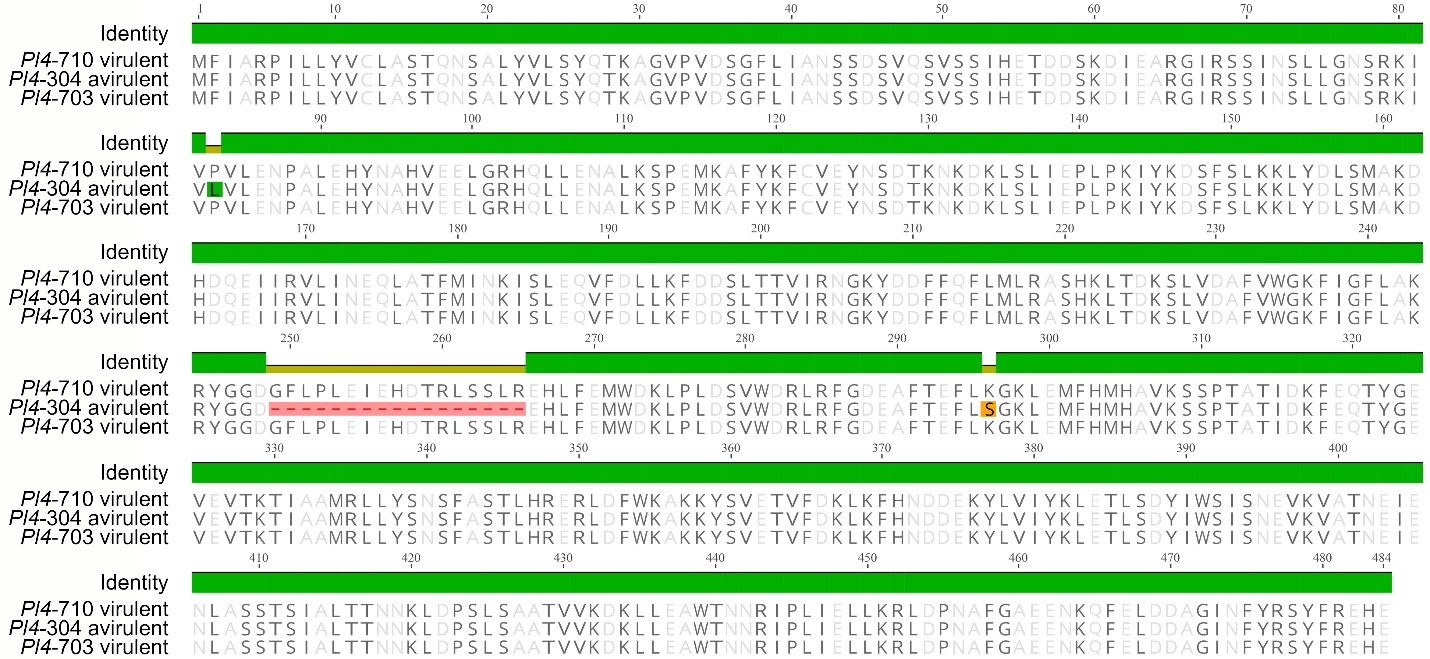

**Fig. S7** Sequence alignment of a candidate effector protein associated with *Pl15* breakdown. Both 710 (Plhal710r2c019g0082461) and 304 (Plhal304r1c010g0039761) alleles confer avirulence towards *Pl15* resistance, they differ only by 5 amino acids at the end of the proteins. The 703 corresponding virulent allele (Plhal703r1c26g0108751) is very divergent and a signal peptide located from 1 to 22 amino acids (position 23-44 on the alignment) is predicted. The 703 allele is found similarly conserved in all virulent pathotypes. It is possible that the 304 and 710 starts of the proteins are mis-predicted, and that they start at methionine 23 allowing the presence of a signal peptide. Divergent amino acids are highlighted in color.

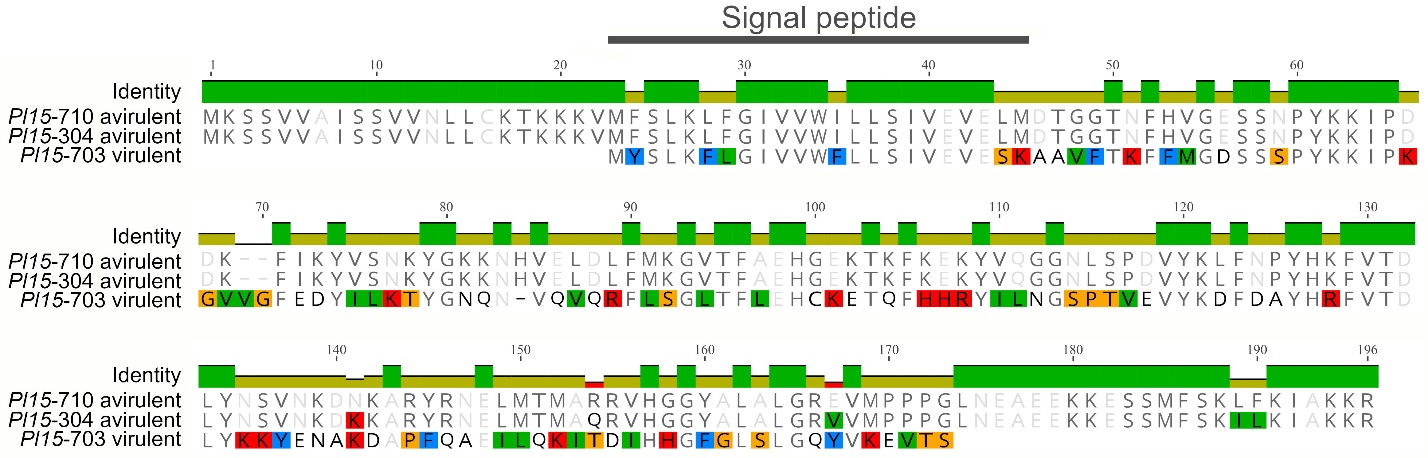

**Table S1** Assembly process summary

**Table S2** Multi-criteria contamination detection

**Table S3** Illumina datasets

**Table S4** *Plasmopara* *halstedii* effector statistics

**Table S5** Secretomes

**Table S6** Effectomes

**Table S7** Summary of regions of interest for resistance breakdowns

**Methods S1** Click here to enter text.

**Notes S1** Click here to enter text.

**Video/Movie S1** Click here to enter text.
